# ARGONAUTE10 is required for cell fate specification and the control of formative cell divisions in the Arabidopsis root meristem

**DOI:** 10.1101/2021.02.05.429893

**Authors:** Nabila El Arbi, Ann-Kathrin Schürholz, Alexei Schiffner, Inés Hidalgo Prados, Friedrich Böhme, Christian Wenzl, Xinai Zhao, Jian Zeng, Jan U. Lohmann, Sebastian Wolf

## Abstract

A key question in plant biology is how oriented cell divisions are integrated with patterning mechanisms to generate organs with adequate cell type allocation. In the root vasculature, a miRNA gradient controls the abundance of HD-ZIP III transcription factors, which in turn control cell fate and spatially restrict vascular cell proliferation to specific cells. Here, we show that a functional miRNA gradient requires an opposing gradient of ARGONAUTE10, which sequesters miRNAs to protect HD-ZIP III transcripts from degradation. In the absence of ARGONAUTE10, xylem precursor cells undergo periclinal divisions that lead to continuous strands of differentiated xylem elements at ectopic positions. Notably, periclinal daughter cells maintain xylem identity even when they are located outside of the xylem axis, resulting in disrupted tissue boundaries. We further demonstrate that ARGONAUTE10 and HD-ZIP IIIs buffer cytokinin signalling to control formative cell divisions, providing a framework for integration of phytohormone and miRNA-mediated patterning.

## Introduction

Plant cells are confined by cell walls that restrict their movement and tether them to neighbouring cells for their entire lifespan. Cells produced by stem cell systems organized in meristems serve as elongation units for morphogenesis, but also acquire specific cell fates based on their position in the organism. Therefore, shape generation in plants depends on oriented cell divisions integrated with developmental patterning and genetically encoded cell fate determination. One of the key evolutionary innovations that allowed plants to dominate the vast majority of terrestrial habitats (Bar-On, et al., 2018), was the development of vascular systems for water and nutrient transport over long distances (Lucas, et al., 2013). The water-conducting xylem tissue of the vasculature is composed of highly specialized cells typically coated by thick secondary cell walls, which fulfil the dual function of stabilizing the xylem elements against the negative pressure of a moving water column and supporting upright growth. Phloem cells, on the other hand, distribute photoassimilates and other molecules throughout the plant body. In the model plant *Arabidopsis thaliana*, it has been firmly established that the precursor cells giving rise to the root vasculature are specified in the embryo (De Rybel, et al., 2016). However, these cells undergo periclinal or radial (i.e. formative) divisions post-embryonically to give rise to additional cell files and thus complete pattern formation. Later in development the procambium, a tissue intervening phloem and xylem, proliferates to give rise to secondary phloem and xylem and thus drives radial growth (Smetana, et al., 2019).

Cell fate determination in plants is mainly based on positional information, rather than cell lineage (Efroni, et al., 2016; Kidner, et al., 2000; Berger, et al., 1998; van den Berg, et al., 1997; van den Berg, et al., 1995). For patterning of vascular cell fates, the interplay of auxin and cytokinin hormone signalling pathways is crucial (Muller and Sheen, 2008; Hamann, et al., 2002; Hardtke and Berleth, 1998). Provascular cells in the centre of the embryonic root are specified by local auxin maxima created through polar auxin transport. Auxin is also involved cell and non-cell autonomously in the establishment of the root quiescent centre, which acts as an organizer for the surrounding stem cells (Schlereth, et al., 2010; Weijers, et al., 2006; Hamann, et al., 1999; Hardtke and Berleth, 1998). This organization of the embryonic root serves as persistent template for the elaboration of the post-embryonic primary root. Formative cell divisions that are initiated directly above the quiescent centre result in a step-wise increase in cell files of the stele, which eventually is constituted by five xylem precursor cells arranged in a single axis, two phloem poles arranged perpendicular to the xylem, the intervening procambium cells, and a peripheral ring of pericycle cells (Figure S1A). High auxin response in xylem precursors causes expression of the basic helix-loop-helix transcription factor TARGET OF MONOPTEROS 5 (TMO5), which together with LONESOME HIGHWAY, another bHLH TF, promotes the biosynthesis of cytokinin (CK) (De Rybel, et al., 2014; Ohashi-Ito, et al., 2014; De Rybel, et al., 2013). However, xylem precursor cells are simultaneously insulated against the effects of CK signalling (Muraro, et al., 2014; Bishopp, et al., 2011; Mahonen, et al., 2006), resulting in non-cell autonomous CK activity in the neighbouring procambial cells. CK signalling in turn channels auxin towards the xylem precursor cells by directly controlling expression and polarity of the PIN class of auxin efflux carriers (Bishopp, et al., 2011; Mahonen, et al., 2000). Importantly, cytokinin signalling is required for formative cell division in the stele, as shown by the reduced number of vascular cell files in plants lacking cytokinin receptors or the ARABIDOPSIS RESPONSE REGULATOR (ARR) transcription factors that mediate CK responses (Argyros, et al., 2008; Ishida, et al., 2008; Yokoyama, et al., 2007; Mahonen, et al., 2006; Mahonen, et al., 2000). At least in part, cytokinin in the stele acts through controlling the expression of a battery of DNA-BINDING WITH ONE FINGER (DOF) transcription factors that promote formative cell divisions (Miyashima, et al., 2019; Smet, et al., 2019). Another group of transcription factors, the class III HOMEODOMAIN LEUCINE-ZIPPER (HD-ZIP III) family proteins, counteract a subset of the DOF factors and inhibit periclinal cell divisions (Miyashima, et al., 2019; Carlsbecker, et al., 2010). The expression of the HD-ZIP III proteins, i.e PHABULOSA (PHB/AtHB-14), PHAVOLUTA (PHV/AtHB-9), CORONA (CNA, AtHB-15), REVOLUTA (REV/AtHB-9), and AtHB-8, is restricted to the centre of the stele by an intricate regulatory mechanism involving mobile miRNAs of the miR165/166 family (Carlsbecker, et al., 2010; Rhoades, et al., 2002). Transcription of the miR165/6 genes in the root occurs specifically in the endodermis that surrounds the vascular cylinder, but the miRNAs move from cell to cell through plasmodesmata and form a gradient with a minimum in the centre of the stele. Presence of miR165/6 leads to the degradation (slicing) of HD-ZIP III transcripts through ARGONAUTE1 (AGO1) (Miyashima, et al., 2011; Carlsbecker, et al., 2010; Kidner and Martienssen, 2004). As a result, HD-ZIP IIIs, which appear to be the main targets of miR165/166, are restricted to the stele centre, even though their promoters can be active in more peripheral cells (Carlsbecker, et al., 2010). Due to the presence of HD-ZIP III proteins, the central cells of the stele generally only divide anticlinally, but not periclinally, and only cells that are in direct contact with the pericycle undergo formative divisions (Miyashima, et al., 2019). HD-ZIP III dosage is also involved in determining xylem tissue type independently of cell division control: high levels of HD-ZIP IIIs in central xylem precursor cells result in the specification of metaxylem with reticulate secondary cell wall thickenings, whereas miR165/6-mediated destruction of HD-ZIP III transcripts in the two cells at the periphery of the xylem axis enables the formation of protoxylem, displaying annular/spiral wall patterning. Thus, control of HD-ZIP III expression by miRNA165/6 (Bao, et al., 2004; Juarez, et al., 2004; Mallory, et al., 2004; Tang, et al., 2003) and AGO1 plays a dual role in restricting formative cell divisions as well as in cell fate determination. The closest relative of AGO1, ARGONAUTE10 (AGO10, also known as ZWILLE and PINHEAD (Moussian, et al., 1998; Endrizzi, et al., 1996; McConnell and Barton, 1995; Jürgens, et al., 1994) has been shown to compete with AGO1 for miRNA165/6 ((Zhu, et al., 2011; Liu, et al., 2009; Mallory, et al., 2009; Lynn, et al., 1999). Since AGO10 does not appear to exhibit slicing activity but rather recruits miRNA-degrading nucleases (Yu, et al., 2017), AGO10 can protect HD-ZIP III transcripts from AGO1-mediated degradation (Liu et al., 2009). AGO10 mutants have been described to display embryonic and post-embryonic defects in the shoot apical meristem (SAM) that can lead to stem cell differentiation and meristem termination (Zhu, et al., 2011; Liu, et al., 2009; Lynn, et al., 1999; Moussian, et al., 1998). However, these severe phenotypes are only observed in a minority of Arabidopsis ecotypes with a restricted geographical distribution (Tucker, et al., 2013; Mallory, et al., 2009; Takeda, et al., 2008). Recently, AGO10 has been revealed to promote meristem formation in leaf axils (Zhang, et al., 2020). Despite these analyses, a role for AGO10 in the root, where miR165/6-mediated regulation of HD-ZIP III transcription factors is instrumental for vascular patterning, has not yet been described. Here, we show that AGO10 is required for the control of formative cell divisions in the centre of the root stele, since its loss leads to ectopic xylem strands, increased procambial cell number, and enhanced root growth. AGO10 acts through protecting HD-ZIP III transcripts from miRNA165/6-mediated degradation, and its promoter activity is graded inversely to the miRNA gradient. Furthermore, AGO10-guarded HD-ZIP III activity buffers cytokinin responses in the root stele to coordinate oriented cell divisions with patterning. Our results demonstrate that AGO10 is required for maintenance of an instructive miRNA165/6 gradient, presumably by adapting steepness of the gradient to the range required by the cellular layout of the root. Presumably as a result of increased vascular cell number, AGO10 mutants outperform the wild type under water-limiting conditions.

## Results

### A mutant with ectopic xylem formation in the root vasculature

Arabidopsis roots display a stereotypical patterning of primary xylem in one axis, with two protoxylem cells at the periphery and three, in rare cases four, metaxylem cells in the centre of the axis (Figure S1A). With the aim to study a gene we hypothesized to be involved in vascular development (*RLP4*, At1g28340), we generated mutant lines in *Arabidopsis thaliana* Col-0 plants using CRISPR/Cas9. We discovered xylem phenotypes in two independent lines that showed frame shift mutations in the target gene, but not in a third homozygous mutant line. After backcrossing to the Col-0 wild type, the phenotype segregated away from the *rlp4* mutation, indicating that the latter was not causative for the observed effects on xylem. We therefore tested whether the phenotype could be derived from Cas9 activity at related loci, but could not reveal any genomic changes at the sites of predicted potential off-targets of the CRISPR guide RNA. Nevertheless, we decided to study the novel xylem mutant, which we termed *the show must go on 1* (*sgo1*), after outcrossing of the Cas9 transgene and backcrossing to the Col-0 wild type. Staining of 6-day-old primary roots with the fluorescent lignin dye basic fuchsin revealed that the recessive (Figure S1B) *sgo1* mutants showed an increased number of cells with lignified protoxylem-like secondary cell wall thickenings (Figure 1A). Whereas wild-type roots showed exclusively two protoxylem cells, *sgo1* roots displayed at least three, in most cases four or five protoxylem-like cells (Figure 1B). These cells were always placed at the periphery of the stele, indicating that general xylem patterning remained intact in the mutant. However, these additional protoxylem cells were frequently placed outside of the xylem axis (Figure 1A). This increase in apparent protoxylem cells did not occur at the expense of metaxylem cells. In contrast, *sgo1* showed slightly increased metaxylem cell number, with infrequent occurrence of lignified cells outside of the xylem axis (Figure 1A, C). Thus, *sgo1* showed increased number of xylem cells (Figure 1D), both within the xylem axis and ectopically in the procambium domain. The basic organisation of xylem patterning, with early differentiating protoxylem at the periphery of the stele and late maturing metaxylem in the centre of the stele, remained intact.

**Figure 1.**
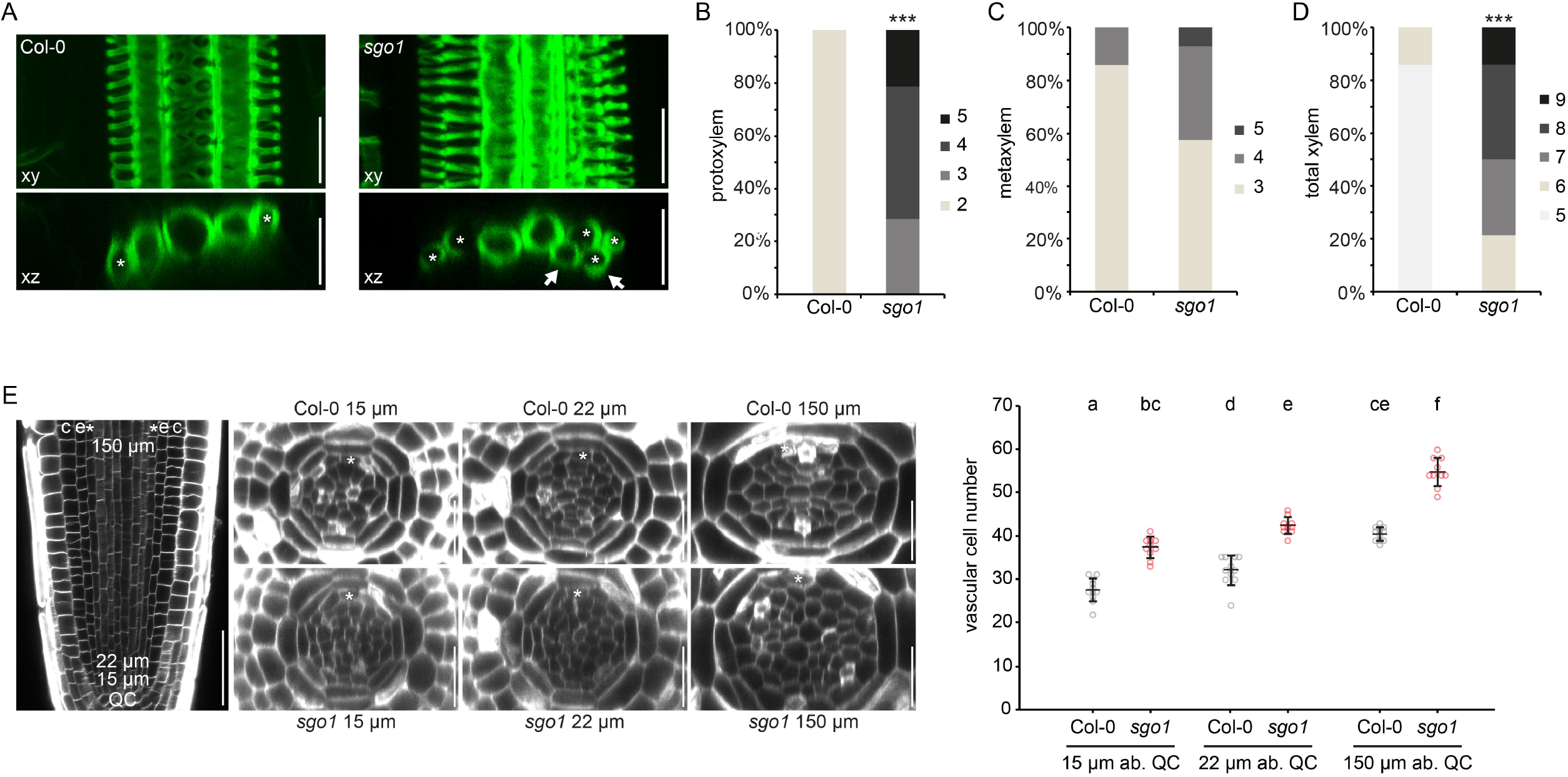
*sgo1* displays ectopic xylem formation and increased formative divisions in the primary root. (A) Lignified xylem cells in Col-0 and *sgo1*. Upper panels are xy projections of a confocal stack, lower panels optical xz sections through the same stack. Asterisks denote cells with protoxylem differentiation, arrows point to ectopic xylem strands. (B-D) Frequency of roots with the indicated number of protoxylem (B), metaxylem (C) or total xylem (D) cells in Col-0 and *sgo1*. Asterisks indicate statistically significant difference from Col-0 based on Mann-Whitney U test (***P<0.001). (E) Quantification of vascular cell number in cross section of confocal stacks at 15 µm, 22 µm, and 150 µm distance from the quiescent centre (QC) cells. Panels on the left show representative Col-0 and *sgo1* meristems. c = cortex, e = endodermis, asterisk = pericycle. Graph depicts means ± s.d. and individual data points (n = 9-11). Letters in graph indicate statistically significant differences based on Tukey’s post-hoc test after one-way ANOVA. Scale bar in left panel = 50 µm, other scale bars = 25 µm.

To assess whether this increased xylem cell number in *sgo1* is due to cell fate changes in the procambium (Holzwart et al., 2018) or rather is the consequence of increased xylem precursor cell division activity in *sgo1* roots, we acquired high resolution 3D stacks of Col-0 and *sgo1* root meristems after tissue clearing and cell wall staining (Ursache, et al., 2018; Musielak, et al., 2015) and generated optical cross sections through the stele. To capture both, the dynamics in the zone of formative cell division and the fully patterned root, we quantified cell number close to the stem cell region, 15 µm and 22 µm above the quiescent centre (QC), and 150 µm above the QC (Figure 1E), where the final stele patterning has been established (Miyashima, et al., 2019). At all positions, *sgo1* cell number was significantly increased compared to the Col-0 wild type, with on average more than 10 additional cells in the mutant at 150 µM above QC (Figure 1E). Thus, *sgo1* shows more formative cell divisions in the root meristem, which could be the cause for the increased number of differentiated xylem cells in the mature part of the root. In wild-type roots, xylem precursor cells in essence never divide periclinally after a QC-proximal formative division of the outer xylem precursor cells resulting in the canonical xylem axis (Miyashima, et al., 2019; Dolan, et al., 1993). Likewise, inner procambial cells without contact to the pericycle remain quiescent and do not contribute any formative divisions until much later in development, when secondary growth is initiated (Miyashima, et al., 2019; Smetana, et al., 2019). To test whether the increased number of xylem precursor cells in *sgo1* could be due to loss of quiescence in xylem precursor and inner procambial cells, we quantified cell division activity in the Col-0 WT and *sgo1* by monitoring incorporation of the thymidine analogue 5-ethynyl-2′-deoxyuridine (EdU). Externally supplied EdU is incorporated during S-phase and can be subsequently coupled to a fluorescent dye by a click-chemistry reaction (Kotogany, et al., 2010). After a 2-hour EdU treatment and subsequent fixation, clearing, and imaging, fluorescent nuclei were dispersed throughout the root meristem (Figure 2A), with *sgo1* displaying slightly higher numbers of S-phase nuclei (Figure 2B). Occasionally, cells with fluorescent nuclei appeared as pairs or clusters, presumably because these cells are the product of recent divisions and thus have a synchronized cell cycle. We then followed a pulse-chase strategy incorporating an EdU-free period of 6 hours following a 40 minute EdU pulse (Figure 2C). We reasoned that this should allow a majority of S-phase cells that acquired EdU during the pulse to proceed through the G2-and M-phases during the chase period. Accordingly, we observed fluorescence occasionally in mitotic structures (Figure 2C, arrow) and predominantly in paired nuclei (Figure 2C, arrowheads). While chance pairings are possible, it can be assumed that a substantial fraction of nuclei pairs are derived from partitioning of replicated DNA to offspring of a mother cell that acquired EdU during the pulse period. Thus, nuclei pairs are not only a read-out for cell division activity, but also indicate cell division orientation (Yin and Tsukaya, 2016). As expected, the majority of fluorescently labelled and paired cells were oriented parallel to the root long axis, indicating an anticlinal division. We focused on pairings periclinal to the root long axis, since these could be derived from recent formative cell divisions. In accordance with the previously observed occurrence of formative cell division exclusively in cells that are in contact with the pericycle (Miyashima, et al., 2019), periclinal pairs of fluorescence nuclei were restricted to positions adjacent to the pericycle in the WT (Figure 2D, F). In sharp contrast, *sgo1* roots showed periclinal pairs throughout the stele, including positions distant from pericycle cells (Figure 2E,F). Notably, the resulting enhanced radial growth of the root did not impede longitudinal root growth; in fact, *sgo1* mutants exhibited longer roots than the wild type (Figure 2G). In summary, SGO1 is required to suppress cell division activity in the centre of the stele, and increased overall cell number in the mutant could be causative for the observed increase in differentiated xylem cells (Fig 1 A-D).

**Figure 2.**
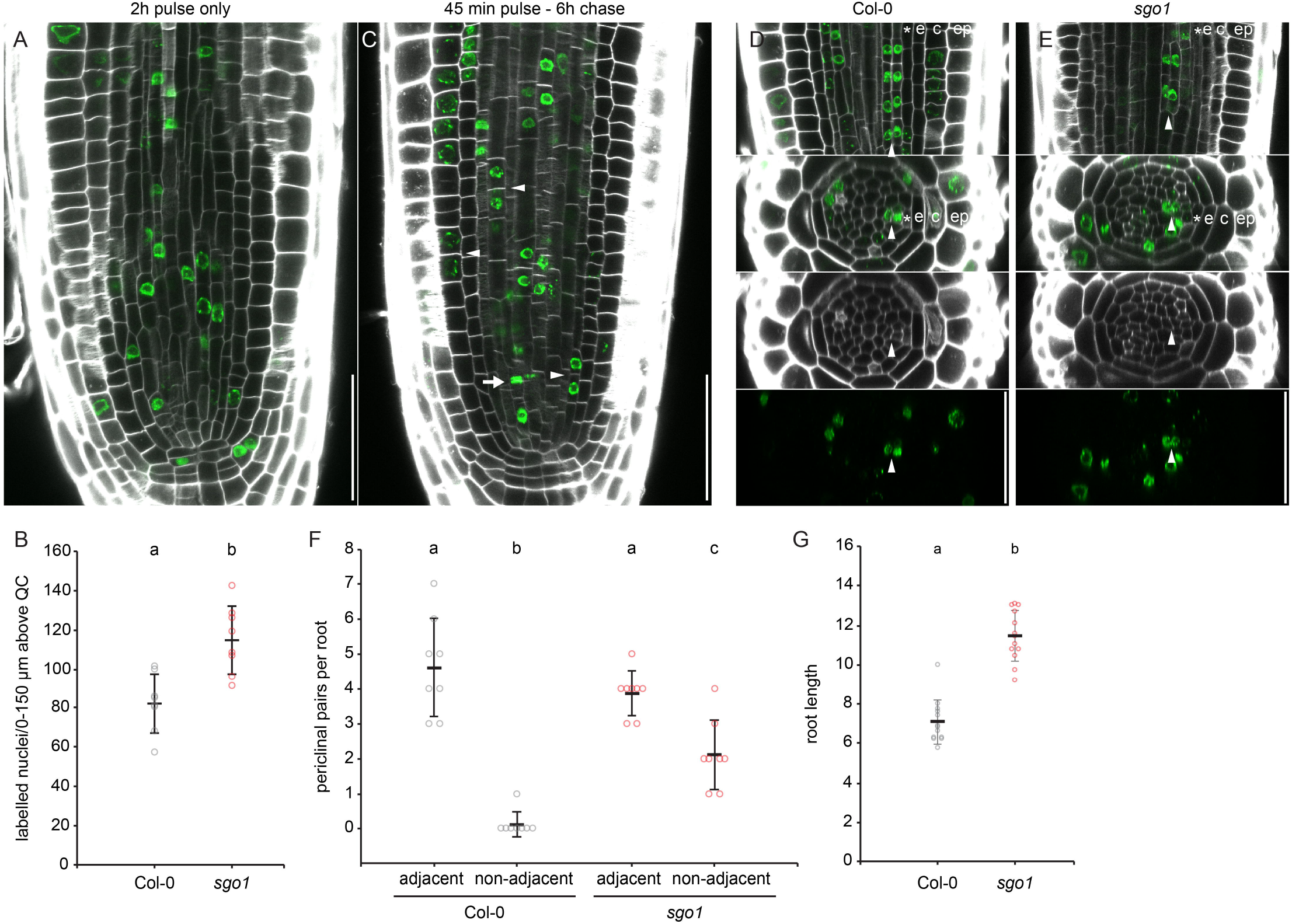
*sgo1* shows enhanced periclinal cell division activity in the root meristem. (A) Exemplary image of a wild-type (Col-0) root with calcofluor white-stained cell walls and Alexa-488-labelled S-phase nuclei after two hours of EdU exposure. (B) Quantification of EdU-labelled nuclei in the vascular cylinder at 0 -150 µm distance from the QC. (C) Exemplary image of a Col-0 root after 45 min of EdU feeding and a 6-hour chase period without EdU. Note labelling of mitotic structures (arrows) and nuclei pairs (arrowheads), likely descendants of cells acquiring EdU during previous S-phase. (D, E) Representative confocal stacks of a Col-0 (D) and *sgo1* (E) meristem showing pairing of nuclei indicative of periclinal cell divisions. Image series shows same periclinal pairs in xy section (upper panel) and xz cross sections (lower three panels) as composite image, cell wall and Alexa-488 channel, respectively. (F) Quantification of periclinal nuclei pairs in the vascular cylinder (excluding pericycle) at 0 -150 µm distance from the QC. Cells adjacent and non-adjacent to the pericycle are discriminated. (G) Root length of Col-0 and *sgo1* seedlings seven days after germination. B, F, and G denote means ± s.d., individual data points are indicated. Letters in graph indicate statistically significant differences based on t-test (B, G) or Tukey’s post-hoc test after one-way ANOVA (F). Scale bars = 50 µm.

### Inner vascular cells ectopically divide and maintain lineage-specific cell identity in *sgo1*

To determine whether *sgo1* roots exhibit an increase in meristematic xylem precursor cells, we crossed the mutant with well-characterized fluorescent marker lines and quantified reporter-positive cells throughout the meristem. The *pAHP6:GFPer* reporter labels protoxylem precursor and adjacent pericycle cells (Mahonen, et al., 2006). We quantified protoxylem precursor cells and observed in the WT background mostly two, occasionally three *pAHP6:GFPer-*positive cells at all three positions in the meristem (15 µm, 22 µm, and 150 µm above QC, Figure 3A, B). While *sgo1* mutants showed similar numbers at the two QC-proximal positions, the *pAHP6:GFPer* expression domain was markedly enlarged at 150 µm above the QC (Figure 3A, B). This enlargement occurred within the xylem axis, but also occasionally included cells outside of the xylem axis, consistent with what was observed in fully differentiated xylem cells. In contrast to p*AHP6:GFPer*, the auxin response marker *DR5v2:YFPer* (Ma, et al., 2019) showed an enlarged expression domain in *sgo1* already at 15 µm above the QC (Figure 3C). It is noteworthy that in the wild type, *DR5v2:YFPer*-derived signal was not strictly correlated with xylem precursor cells directly above the QC but showed quite variable and widespread expression, which stabilized and constricted to the five xylem precursor cells as patterning was completed at 150 µm distance from the QC (Figure 3C). In *sgo1*, the *DR5v2:YFPer* expression pattern remained variable at all positions and was markedly expanded compared to the wild-type background (Figure 3C). As was observed with p*AHP6:GFPer*, enlargement of marker expression domain was due to an increased number of cells in the xylem axis as well as to reporter-positive cells outside the axis. In a next step, we crossed *sgo1* with a marker line driving nuclear GFP expression under control of the TMO5 promoter to quantify the number of cells with xylem identity in the meristem (Schlereth, et al., 2010). TMO5 expression is largely controlled by auxin, however the reporter exhibits a more restricted activity compared to *DR5v2:YFPer*. In most wild-type roots, *TMO5:NLS-3xGFP* was confined to five cells (Figure 3D), consistent with faithful representation of xylem precursor cells. In sharp contrast, the number of *TMO5:NLS-3xGFP*-positive cells in *sgo1* roots was substantially increased throughout the meristem (Figure 3D). Quantification of pTMO5-positive cell files that were the apparent result of ectopic xylem precursor cell divisions, revealed on average more than three of those cell files in *sgo1* (Figure 3E). Consistent with expectations, ectopic xylem precursor cell files were virtually absent in the wild-type background. Likewise, and in contrast to the Col-0 wild type, *sgo1* roots frequently showed additional cell files resulting from formative divisions in inner procambium cells directly adjacent to *pTMO5:NLS-3xGFP*-positive cells (Figure 3F). It is noteworthy that the increased formative division in *sgo1* did not occur randomly but led to continuous strands of TMO5-positive precursor cells, even when the extra strands appeared displaced from the xylem axis. However, the length of these ectopic strands varied. We could observe ectopic TMO5-positive cell files that spanned the entire meristem (Figure S2) as well as the initiation of ectopic strands (Figure 3E, arrow) and beginning displacement form the meristem (Figure 3F, arrows). This is consistent with our observations in the mature part of the root, where additional xylem files are also largely continuous (Figure 1). In summary, SGO1 is involved in the control of stele cell number and xylem proliferation.

**Figure 3.**
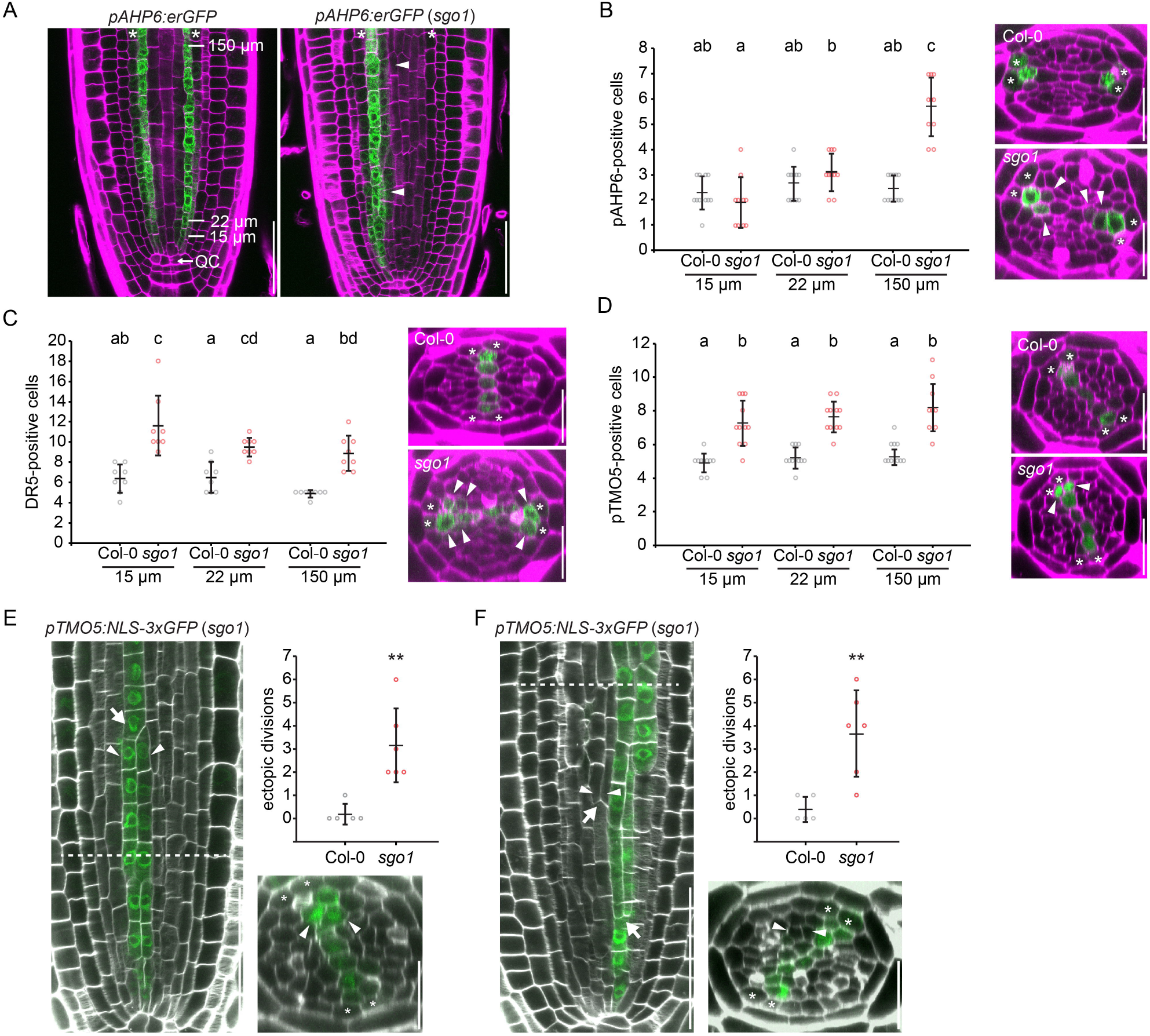
Inner vascular cells ectopically divide and maintain lineage-specific cell identity in *sgo1*. (A) Expression of the *pAHP6:GFPer* marker is expanded inward in *sgo1*, while expression in pericycle cells is reduced. Asterisks = pericycle. (B) Quantification of *pAHP6:GFPer*-positive xylem cells in cross sections of confocal stacks of WT and *sgo1* at the indicated distances from the QC. (C) Quantification of *pDR5v2:YFPer*-positive xylem cells in cross sections of confocal stacks of Col-0 and *sgo1*. (D) Quantification of *pTMO5:NLS-3xGFP*-positive xylem cells in cross sections of confocal stacks of Col-0 and *sgo1*. (E) *pTMO5:NLS-3xGFP*-positive xylem precursor cells ectopically divide in *sgo1* but not Col-0. Arrow points to initiation of a continuous ectopic cell file. (F) Inner procambial cells adjacent to *pTMO5:NLS-3xGFP*-labelled xylem precursor cells ectopically divide in *sgo1*. Arrow points to a continuous ectopic cell file being displaced from the meristem. Dashed lines indicate position of xz section. Graphs in (B-D) depict means ± s.d., individual data points are indicated. Letters indicate statistically significant differences based on Tukey’s post-hoc test after one-way ANOVA. Graphs in (E) and (F) display mean of ectopic divisions per root (0-150 µm distance from QC) ± s.d., individual data points are indicated. Asterisks in graphs indicate statistically significant differences based on student’s t-test (**P<0.01). Asterisks in micrographs denote pericycle cells at xylem poles, arrowheads indicate ectopic marker expression (A-E) or ectopic periclinal division (F). Scale bar in xy sections = 50 µm, in xy sections = 25 µm.

### *SGO1* encodes AGO10

Through a combination of bulked segregant analysis and whole genome sequencing, we identified a single nucleotide polymorphism (G1333A counting from ATG) in the *AGO10* coding sequence in *sgo1*, resulting in an amino acid substitution from E to K at position 445 in the AGO10 protein. To determine whether mutation of AGO10 was causative for the *sgo1* phenotype, we analysed the AGO10 loss-of-function mutant *zll-3* (Moussian, et al., 1998) in the Landsberg erecta (L*er*) background. *zll-3* roots displayed additional differentiated xylem files as well as increased number of meristematic cell files, very similar to the *sgo1* mutant (Figure S3A-C). Likewise, roots of the *ago10-1* mutant (Takeda, et al., 2008) which is a T-DNA insertion line in the Col-0 background, showed a meristematic cell number phenotype indistinguishable from *sgo1*, suggesting that the *ago10* SNP in *sgo1* is the causative mutation (Figure 4A). Consistent with this, F1 trans-heterozygotes resulting from a cross of *sgo1* with *zll-3* showed the *sgo1*/*zll-3* xylem phenotype, demonstrating that *sgo1* is a novel AGO10 loss-of-function mutant (Figure S3D).

**Figure 4.**
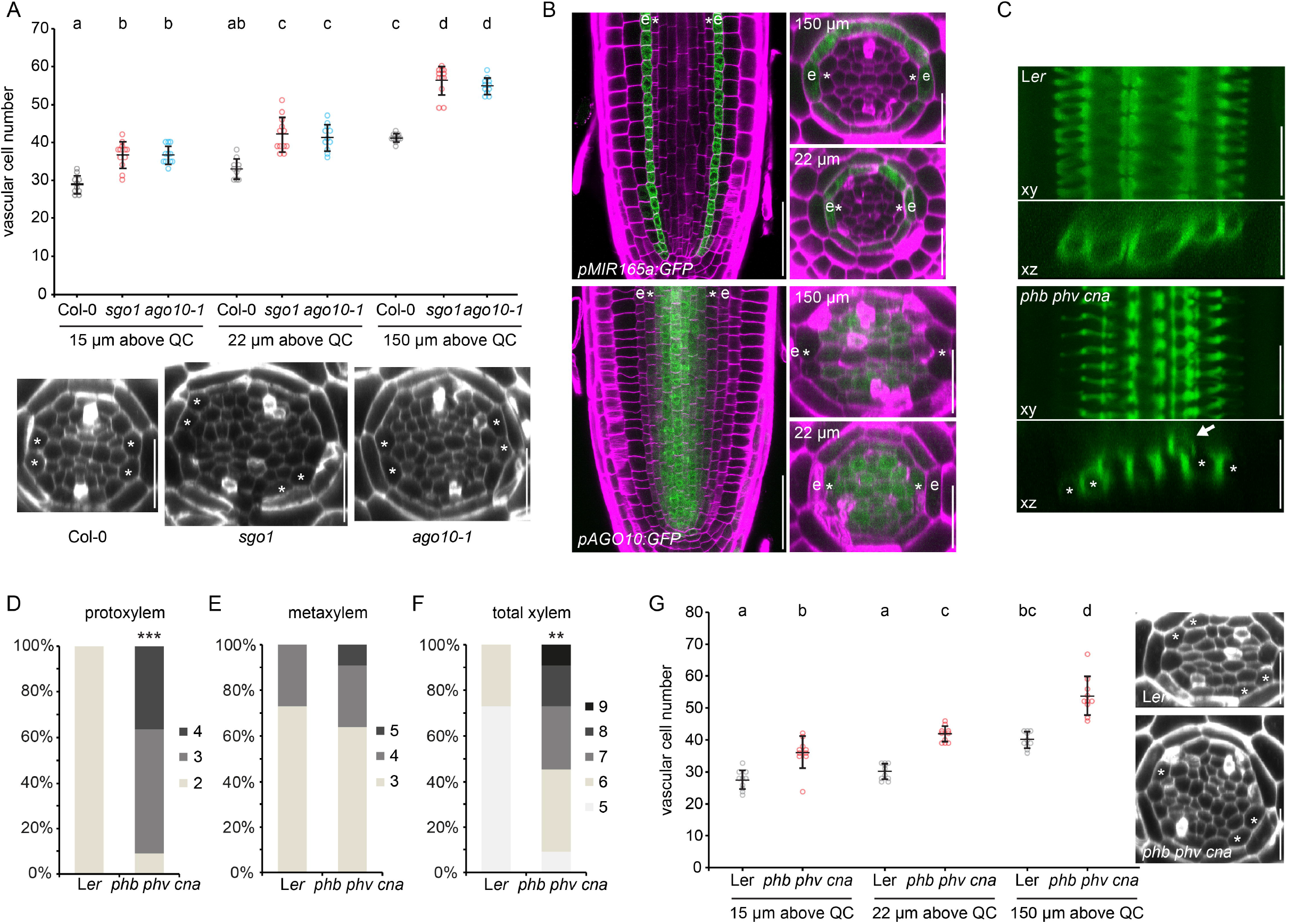
AGO10 is required for HD-ZIP III-mediated vascular patterning. (A) Quantification of vascular cell number in cross section of confocal stacks. Graph depicts means ± s.d. and individual data (n = 10-12). Letters in graph indicate statistically significant differences based on Tukey’s post-hoc test after one-way ANOVA. (B) Expression of *pMIR165a:GFP* and *pAGO10:GFP* reporters in the root meristem. Longitudinal xy sections as well as xz sections from confocal stacks are displayed. e = endodermis, asterisks = pericycle. (C) Staining of lignified xylem cells. Upper panels are xy projections of a confocal stack, lower panels depict xz sections through the same stack. (D-F) Frequency of roots with the indicated number of protoxylem (D), metaxylem (E) or total xylem (F) cells in L*er* and *phb phv cna*. Asterisks indicate statistically significant difference based on Mann-Whitney U test (***P<0.001, **P<0.01). (G) Quantification of vascular cell number in cross sections of confocal stacks. Graph depicts means ± s.d. and individual data points. Letters indicate statistically significant differences based on Tukey’s post-hoc test after one-way ANOVA. Asterisks = pericycle.

### AGO10 is required for HD-ZIP III function in the root vasculature

AGO10 has been described to specifically bind miR165/166 and efficiently compete with AGO1 for these miRNAs due to a higher affinity (Zhu, et al., 2011). As it shows no slicing activity, AGO10 is assumed to negatively regulate AGO1-mediated degradation of transcripts of HD-ZIP III transcription factors, the targets of miR165/166. In the root, the promoters of the miR165/166 genes are exclusively active in the endodermis (Miyashima, et al., 2011; Carlsbecker, et al., 2010) (Figure 4B). This tissue specific expression, together with (regulated) movement through plasmodesmata (Skopelitis, et al., 2018; Vaten, et al., 2011) establishes a miRNA gradient with peak abundance in the endodermis and periphery of the stele, and a minimum in and around the metaxylem cells. Accordingly, HD-ZIP III expression, with the possible exception of REV, is mostly confined to the inner vascular cells (Carlsbecker, et al., 2010). A reporter line expressing nuclear envelope-localized GFP under control of the AGO10 promoter-driven reporter fluorescence (*pAGO10:GFP*) (Palovaara, et al., 2017)) suggested a gradient of AGO10 transcription inverse to miR165/166 abundance with very low signal in the endodermis and progressively increasing reporter fluorescence towards the centre of the stele. Interestingly, the *pAGO10:GFP* reporter was active in all inner vascular cells in the region directly above the QC, but reporter fluorescence markedly decreased in xylem precursor cells towards the shoot (Figure 4B, Figure S4A). Thus, inverse gradients of miR165/6 and AGO10 might be cooperating to control HD-ZIP III expression in the root stele. Consistent with the proposed role of AGO10 in protecting HD-ZIP III transcripts against slicing, quantitative real time PCR analysis demonstrated a decrease in the abundance of HD-ZIP III transcripts in *sgo1* (Figure S4B). Moreover, a *phb phv cna* mutant (Prigge, et al., 2005) showed root phenotypes remarkably similar to *ago10* mutants (Figure 4C-G): supernumerary and ectopic differentiated xylem cells as well as increased meristematic stele cell number essentially phenocopied *ago10* mutants, again without compromising root growth (Figure S4C), indicating that AGO10 indeed mainly acts through controlling HD-ZIP IIIs. In agreement with this, sequestration of miR165/6 by expression of a target mimic (*35S:STTM165/6*) (Yan, et al., 2012) alleviated the *sgo1* phenotype (Figure S4D). Taken together, our results suggest that in the absence of AGO10, the miR165/166 gradient is too shallow to allow for sufficient HD-ZIP III expression, consistent with a dual function of AGO10 in sequestering miR165/6 as well as its degradation.

### AGO10 and HD-ZIP III proteins buffer vascular cytokinin responses

It has been shown that cytokinin is essential for formative divisions in the root vasculature (De Rybel, et al., 2014; Ohashi-Ito, et al., 2014; Mahonen, et al., 2006; Mahonen, et al., 2000). We therefore investigated whether the CK response might be affected in *ago10* mutants using a CK response reporter (Zurcher, et al., 2013). Indeed, *pTCSn:2xVenus-NLS* showed an enlarged expression domain in the *sgo1* mutant compared to the wild type as well as increased reporter fluorescence in the stele. In particular, *pTCSn:2xVenus-NLS*-derived fluorescence was observed throughout the procambial tissue in the root meristem, whereas reporter activity in wild-type roots sharply decreased a few cells above the QC (Figure 5A). The addition of the synthetic cytokinin 6-benzyl adenine (BAP) increased the intensity of reporter-derived fluorescence in both wild type and *sgo1* backgrounds but it did not alter the sharp decrease towards the more mature parts of the wild-type roots (Figure 5A). This strongly suggests that the vascular cytokinin response is buffered in the transition zone of the root and that this buffering mechanisms depends on *AGO10*, and, presumably, on HD-ZIP III TFs. Interestingly, expression of *AGO10* itself is responsive to CKs, since *pAGO10-GFP* activity displayed an increase in fluorescence intensity and expression domain, including the xylem precursor cells, at 150 µm above the QC upon BAP treatment (Figure 5B). We then tested the response of Col-0, *sgo1*, L*er*, and *phb phv cna* roots to increased CK signalling with respect to formative cell divisions in the root stele by quantifying vascular cell number under different concentrations of BAP at 150 µm above the QC. Whereas both the Col-0 and L*er* wild types did not show a significant change in vascular cell number in response to up to 10 µM BAP, *sgo1* and *phb phv cna* showed a dramatic increase of formative cell divisions (Figure 5C). In extreme cases, this could result in roots with more than 100 cells, as opposed the the average of ∼40 cells observed in the corresponding wild-type roots. This result shows that buffering of the promoting effect of CK signalling on formative cell divisions in the stele is mediated by the AGO10-HD-ZIP III module.

**Figure 5.**
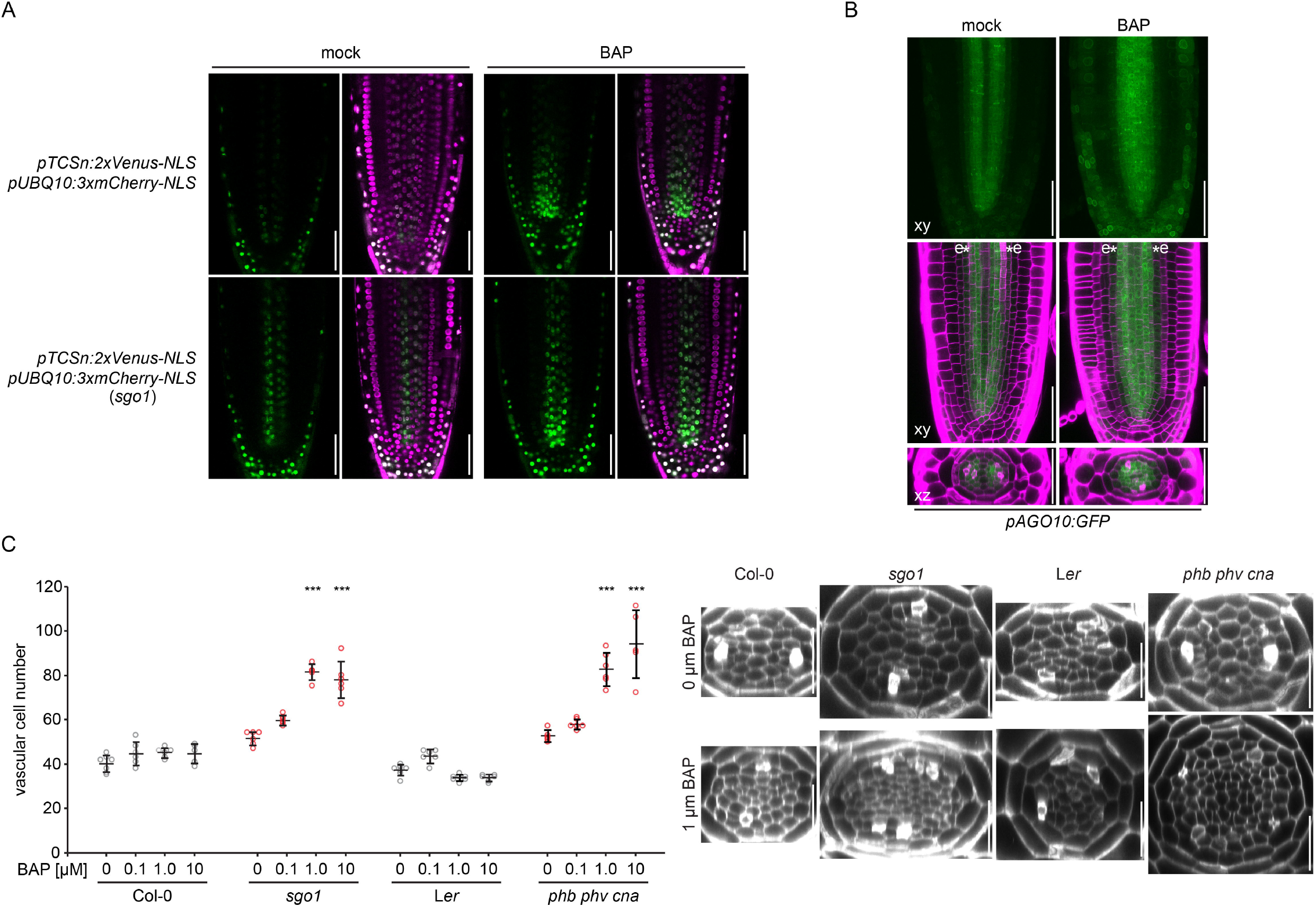
AGO10 and HD-ZIP III proteins buffer vascular cytokinin responses. (A) Confocal images of *pTCSn:2xVenus-NLS pUB10:3xmCherry-NLS* marker line after mock treatment or after growth on medium supplemented with 0.1 µM BAP (upper panels). Lower panels show the same marker line in the *sgo1* background. (B) *pAGO10:GFP* expression as displayed in Fig 3 after mock treatment or after growth on 0.1 µM BAP. (C) Quantification of vascular cell files of Col-0, *sgo1*, L*er*, and *phb phv cna* roots in response to increasing concentrations of BAP. Graph depicts means ± s.d. and individual data points. Asterisks indicate statistically significant differences from the respective control (0 µM BAP) within the same genotype based on Tukey’s post-hoc test after one-way ANOVA (***P<0.001). Scale bars in (A) and (B) = 50 µm, in (C) = 25 µm.

Finally, we tested whether AGO10-mediated control of root vascular architecture affects plant performance under environmental stress conditions. As an increase in xylem cell number is associated with plant adaption to water stress (Ramachandran, et al., 2020), we compared the growth of WT and *sgo1* plants on medium simulating water limitation by addition of polyethylene glycol (PEG) (Verslues and Bray, 2006). While vascular cell number was reduced under these conditions in both WT and *sgo1* (Figure S5A), root growth of *sgo1* plants was more resistant to water-limiting conditions. This was the case when the plants were germinated and grown on PEG-containing medium (Figure S5B), as well as after transfer to PEG medium following initial cultivation on control medium (Figure S5C).

## Discussion

Plants rely on integration of oriented cell division with developmental patterning of cell identities for the generation of functional organs and tissues. Correct cell fate determination is particularly important in the vasculature to ensure connectivity of the conductive tissues that transport water and nutrients throughout the plant body. In this study, we demonstrate that AGO10 is required for the control of formative cell divisions in the root vasculature and is an essential component of miR165/6-mediated patterning. It is firmly established that AGO10 competes with AGO1 for miR165/6 (Zhu, et al., 2011). AGO1 and AGO10 are 78% identical in the miRNA-binding PAZ and target cleaving PIWI domain, and both require 5’ uridine for efficient miRNA loading (Zhu, et al., 2011). However, whereas AGO1 engages with many species of small RNAs to mediate target slicing, AGO10 appears to lack cleaving activity and exclusively associate with the miR165/6 family. Initially, AGO10 mutants were identified in genetic screens performed in the L*er* background due to their strong SAM termination phenotype (McConnell and Barton, 1995; Jürgens, et al., 1994). However, this phenotype is not observed in the most commonly used Col-0 phenotype (Mallory, et al., 2009; Takeda, et al., 2008) and seems to be restricted to Arabidopsis isolates from a narrow geographical distribution (Tucker, et al., 2013). The mutation in *sgo1* leads to a substitution of E445 by lysine, located towards the end of the 3’ miRNA-binding PAZ domain. This residue is also mutated in the L*er zll-7* allele (E445A), exhibiting a phenotype indiscernible from the strong *zll-3* (L*er*) mutant, which fails to maintain a SAM in more than 85% of seedlings (Moussian, et al., 1998). *sgo1, zll-3*, and the T-DNA insertion allele *ago10-1*, which shows dramatically reduced levels of AGO10 transcript (Mallory, et al., 2009; Takeda, et al., 2008), were phenotypically indistinguishable in terms of vascular patterning. Thus, in contrast to the ecotype-specific contribution of AGO10 to shoot apical meristem maintenance, its role in controlling formative divisions in the root seems to be more general. Gene regulation by small RNAs gradients is thought to accomplish reduction of gene expression noise (Schmiedel, et al., 2015), sharpening of expression boundaries (Levine, et al., 2007), or even randomization of cell fate determination (Plavskin, et al., 2016). In the root vasculature, patterning is dominated by the auxin-cytokinin antagonism, which partitions the root in two symmetrical cytokinin response domains (the procambium) and an auxin response domain (the xylem precursors) separating the former. The miR165/6 gradient has its source in the endodermis, outside of these fields, and thus confers the unique property of regulating its targets independent of the auxin-cytokinin interplay. Thus, in the case of HD-ZIP IIIs, expression can be spatially restricted to the centre, but encompass cells of both the auxin and the cytokinin response domain (Carlsbecker, et al., 2010).

HD-ZIP III transcription factors in the stele have a dual role in the repression of formative cell divisions and the dose-dependent determination of meta- and protoxylem cell fate: low expression of HD-ZIP IIIs in the peripheral two xylem precursor cells leads to protoxylem formation, whereas a high HD-ZIP III dose allows for metaxylem differentiation in the centre (Carlsbecker, et al., 2010). Our results showing that *ago10* mutants are indistinguishable from higher order *hd-zip III* mutants would suggest that, in the absence of AGO10, miRNA165/6 can spread further from the endodermal source into the stele, curtailing HD-ZIP III expression. AGO10 leads to degradation of miRNAs by recruiting SDN1 and SDN2 (Yu, et al., 2017), and miR165/6 levels are increased in the absence of AGO10 (Liu, et al., 2009), suggesting that to maintain an instructive miRNA gradient, AGO10 does not only sequester miR165/6 but also triggers its degradation. This would also suggest that, if unchecked by AGO10, miR165/6 might be capable of moving further than previously assumed (Benkovics and Timmermans, 2014) consistent with the experimentally determined range of artificial miRNAs of 10-15 cells (de Felippes, et al., 2011). In summary, AGO10 is an essential regulator of root vascular patterning, most likely through expanding the amplitude of the miR165/6 gradient. It will be interesting to see whether comparable “expander” components are general features of gradient decoding.

Notably, recent results point towards a role of AGO proteins in the active removal of other small RNAs (Devers, et al., 2020). Furthermore, modelling of vascular patterning has predicted that diffusion of miRNA165/6 across cell boundaries is insufficient for correct vascular patterning, and the existence of a mechanism for removal of the miRNA has been hypothesized (Muraro, et al., 2014). Based on our results, we propose that AGO10 might provide this predicted function. One of the key features of gradients formed by mobile small RNAs is that threshold-based readout allows for flexible determination of gene expression boundaries. Remarkably, in leaves, miR165/6 gradient readout does not depend on feedback from an underlying polarity network or additional pre-patterned factors, as a binary on-off interpretation of target gene expression was maintained even if the source of the gradient was altered (Skopelitis, et al., 2017). In contrast to leaves, where the miR165/6-HD-ZIP III module is involved in adaxial-abaxial patterning (Nogueira, et al., 2007; Juarez, et al., 2004), miR165/6 in the root generates an inversely graded expression of HD-ZIP III expression (Miyashima, et al., 2011), suggesting that readout of small RNA gradients differs between leafs and stele. However, one of the consequences of HD-ZIP III expression, suppression of formative cell divisions, occurs with a sharp boundary between cells adjacent to pericycle cells and those further towards the stele centre (Miyashima, et al., 2019). Interestingly, gradient readout is sensitive to the miRNA levels at the source (Skopelitis, et al., 2017; Miyashima, et al., 2011). It is thus conceivable that the interplay of AGO10 and the miRNA165/6 gradient potentiates the flexibility and tuning capacity of patterning. In light of this, it is tempting to speculate that modulation of AGO10 expression might be involved in developmental response to environmental cues. Upon drought stress, roots deploy an increased number of xylem strands, potentially to maximize water transport. Recent studies show that this developmental adaptation is mediated by ABA signalling, which leads to increased miR165/6 expression, and thus a reduction in HD-ZIP III transcript abundance and an increase in protoxylem differentiation (Bloch, et al., 2019; Ramachandran, et al., 2018). In addition, Bloch et al., have shown a decrease in *AGO10* expression after ABA treatment (Bloch, et al., 2019). Here, we show that lesion of AGO10 leads to better performance under water-limiting conditions, possibly due to increased conductance of the xylem. Consistent with the notion that expression of AGO10 can be altered to control patterning, a recent study has found that multiple pathways converge on the AGO10 promoter (Zhang, et al., 2020). In this context, AGO10 acts mainly through REV, which has been shown to counteract the function of other HD-ZIP IIIs in certain situations (Prigge, et al., 2005). Zhang et al., observed only a mild influence of CK on AGO10 expression, despite the presence of CK response elements in the promoter. These discrepancies suggest that alternative wiring of the miRNA-AGO10-HD-ZIP III module may provide flexibility for various developmental contexts.

A key remaining question is how the miR165/6-AGO10-HD-ZIP III module controls formative cell divisions. A recent study has revealed that HD-ZIP IIIs antagonize DOF transcription factors, which promote formative divisions, by repressing DOF transcription and cell to cell movement (Miyashima, et al., 2019). In addition to these direct effects, our results demonstrating that HD-ZIP IIIs buffer cytokinin responses suggest also an indirect level of control, as expression of the PEAR class of DOF TFs is induced by CK signalling (Miyashima, et al., 2019; Smet, et al., 2019) and DOF promoters can be targets of ARRs (Smet, et al., 2019; Xie, et al., 2018; Zubo, et al., 2017). Thus, increase in PEAR expression due to loss of HD-ZIP IIIs might at least partially explain the increase in formative cell division in *sgo1* and *phb phv cna*. Interestingly, PHB has been shown to suppress B-type ARR activity (Sebastian, et al., 2015), although it should be noted that the interactions between CKs and HD-ZIP III are complex and presumably context-dependent (Sebastian, et al., 2015; Dello Ioio, et al., 2012). Both pathways also converge on xylem patterning, as CK application suppresses protoxylem specification and CK mutants show protoxylem in place of metaxylem (Argyros, et al., 2008; Yokoyama, et al., 2007; Mahonen, et al., 2006; Mahonen, et al., 2000), whereas HD-ZIP IIIs gain of function alleles show metaxylem in place of protoxylem (Carlsbecker, et al., 2010). External application of CK leads to a decrease in root length and meristem size. However, *sgo1* and *phb phv cna* roots are longer than WT, despite the increase in CK signalling response. It is well known that individual cell types and tissues respond differentially to CK inputs. For example, CK signalling promotes differentiation in the outer tissues of the root but suppresses xylem differentiation (Dello Ioio, et al., 2007; Mahonen, et al., 2006). Thus, even though CK is essential for correct vascular patterning by promoting formative divisions (De Rybel, et al., 2014; Ohashi-Ito, et al., 2014; Mahonen, et al., 2006; Mahonen, et al., 2000), cell type specific buffering of CK responses by HD-ZIP III might be required to coordinate CK responses in different tissues. In this scenario, the observed CK-responsiveness of AGO10 expression might further enhance buffering of CK responses.

In summary, we demonstrate that AGO10 is required for controlling lateral growth and cell fate determination in the root vascular tissue. Our results indicate that AGO10 is an essential component of the miR165/6 –HD-ZIP III module, enabling the establishment of an instructive miRNA gradient. We speculate that this intricate non-cell autonomous regulatory circuitry simultaneously provides robustness and flexibility to adapt vascular patterning to both developmental and environmental cues.

## Acknowledgements

The authors would like to thank Dolf Weijers, Bert De Rybel, Ari Pekka Mähönen, and Thomas Greb for sharing materials. Furthermore, the authors are thankful to Ingrid Lohmann for providing microscope access, and to Rosa Lozano-Durán for critical reading of the manuscript and discussions. Research was supported by the German Research Foundation (DFG) with grants WO 1660/2 and WO 1660/6 to S.W.

## Author contributions

N.E .A., A.-K. S. J.U.L., and S.W. designed research, N.E .A., A.-K. S, A.S., I.H.P., F.B. C.W., X.Z., J.Z., and S.W. performed experiments and analysed data. S.W. supervised research and wrote the manuscript.

## Materials and Methods

### Plant material and growth conditions

Plant material used in this study is described in the resource table. Seeds were surface-sterilized with 1.2% NaOCl in 70% ethanol for five minutes, followed by two rinses in absolute ethanol and air drying under the sterile bench. If not indicated otherwise, plants were grown on half-strength MS medium supplemented with 1% sucrose and 0.9 % phytoagar. If appropriate, 6-benzyl adenine was added after autoclaving. For growth under simulated water deficit, half strength MS plates were infused with PEG as described previously (Verslues and Bray, 2006). Briefly, 25 ml of half strength MS medium containing 1.5 % plant agar were poured into square 12 cm petri dishes and allowed to solidify. This was overlaid with 37.5 ml of liquid half-strength MS medium containing 0, 250, or 400 g/L of PEG 8000 added after autoclaving. Plates were allowed to dry for three days and excess solution was poured off. On plates overlaid with 400 g/L PEG 8000, seeds failed to germinate. For transfer experiments, seedlings were grown on half-strength MS medium without sucrose and transferred to PEG plates after five days of growth.

### Generation of transgenic plants

For the *pTCSn:2xmVenus-NLS pUBQ10:3xmCherry-NLS* line we first created transgenic plants harbouring construct pCW066 (pUBQ10:3xmCherry-NLS) that were then transformed with construct pCW178 (pTCSn:2xVenus-NLS). Constructs were generated using GreenGate cloning as previously described (Lampropoulos, et al., 2013). Modules and oligo sequences can be found in the Supplemental Tables.

### Identification of *sgo1*

To identify the mutation causal for the sgo1 phenotype we combined bulked segregant analysis with next generation sequencing. We first created a mapping population by crossing *sgo1* to the L*er* wildtype. We scored the xylem phenotype of 200 F2 plants by light microscopy, pooled all the plants with *sgo1* phenotype and analysed with a set PCR-based markers (∼5 per chromosome) for Col-0/Ler polymorphisms. We found a clear bias for Col-0 on the bottom of chromosome 5. We then collected 50 plants each with WT and *sgo1* phenotype from an F2 population resulting from a backcross of *sgo1* to Col-0, prepared genomic DNA and sequenced the two pooled samples together with an *sgo1* sample.

Genomic DNA for whole genome sequencing was prepared using CTAB. The samples were ground under liquid nitrogen using pistil and mortar and 70 – 100 mg plant material was resuspended in 600 μL CTAB buffer (2 % CTAB, 1 % PVP 4000, 1.4 M NaCl, 100 mM Tris-HCl, pH 8, 20 mM EDTA, pH 8). After a one-hour incubation at 65 °C, samples were cooled down to room temperature and 1 μL RNaseA (1 mg/mL) was added. Samples were incubated for one hour at 37 °C. 60 μL CHCl3 were added, gently mixed by inverting the tubes and centrifuged at 5000 g for 10 min at RT. The polar phase (∼ 500 μL) were transferred to a new reaction tube, and nucleic acids were precipitated by adding 2.5 volumes of 100 % ethanol (−20 °C) and incubation for 30 min at – 20 °C. Afterwards, precipitated DNA was pelleted at 11.000 g for 10 min at 4 °C. The supernatant was discarded and 500 μL of 70 % ethanol (−20 °C) were added to wash the precipitated DNA, followed by centrifugation at 11,000 g for 10 min at 4 °C. The supernatant was discarded and the DNA pellet was dried at 55 °C until residual ethanol had evaporated. The DNA pellet was resuspended in 50 μL H2O. Raw sequencing reads were processed with Trimgalore for adapter removal and quality filtering (https://www.bioinformatics.babraham.ac.uk/projects/trim_galore/; (Martin, 2011) and aligned to the TAIR10 reference genome using bwa (Li and Durbin, 2009). After BAMconversion and indexing with SAMtools (Li, et al., 2009), SNPs in the sgo1 alignment were called using bcftools (Li, et al., 2009). The three alignments (*sgo1, sgo1*xCol-0 BCF2 with wild-type phenotype, *sgo1*xCol-0 BCF2 *sgo1* phenotype) were then compared and candidate SNPs identified in the region of interest on chromosome five determined by bulked segregant analysis. Only the SNPs in *AGO10*/*ZWILLE* co-segregated with the *sgo1* phenotype.

### Expression analysis

Seeds were surface-sterilized with chloride gas, sown on nylon mesh placed on half-strength MS plates containing 0.8% Phyto agar, vernalised for 2 days at 4 °C in the dark and vertically grown under long-day conditions (16 h day/8 h night at 22 °C). About 5 cm long roots were cut by blade from 7-day-old seedlings, transferred to protoplast isolation solution (20 mM MES, 400 mM Mannitol, 20 mM KCl, 1.5% Cellulase, 0.4% Macerozyme, 10 mM CaCl2) for 15 min to release the root tip from the more mature part of the root at the transition to the elongation zone. Total RNA including miRNA were isolated from root tips with the miRNeasy Mini Kit (QIAGEN, Cat No. 217004) following the standard protocol. Transcript estimation was performed with by quantitative reverse-transcription polymerase chain reaction (qRT-PCR). RNA extractions of three biological replicates per genotype were DnaseI (Invitrogen)-treated to eliminate any residual DNA contamination. A total of 1 µg RNA per sample was reverse transcribed to cDNA according to the manufacturer’s instructions (AMV Reverse Transcriptase Native, EURX) using an oligodT primer. QRT-PCRs were run in a Rotor-Gene Q 2plex (Qiagen) cycler on diluted cDNA with four technical replicates and quantified in reference to

ACT2. Sybr Green I nucleic acid gel stain (Sigma-Aldrich) was used for detection. Primer efficiency was determined from cDNA serial dilution series’. Reactions were amplified at 95°C initial denaturation for 6min, then looped through 95°C for 30sec, 59°C and 72°C for 30sec or 10sec for miRNA166 for 40 cycles. Melt curves were obtained from 55 to 95°C incrementing 1°C per step. The data were analysed with the 75 Rotor-Gene Q 2plex software and evaluated according to (Muller, et al., 2002). Statistics were performed according to (Rieu and Powers, 2009).

### Genotyping

The presence of the *sgo1* SNP in AGO10 was assessed with a derived cleaved amplified polymporphic sequence (dCAPS) marker (see table X for oligo sequences). Mutant, but not wild-type amplicons can be cleaved with HindIII, resulting in fragments of 137 bp and 31 bp that were resolved on a 3 % agarose gel. Genomic DNA for genotyping was extracted by grinding ∼100 mg of leaf tissue in a homogenizer (Retsch mill, QIAGEN) for 30 s and 30 rpm. After addition of 250 μL of gDNA extraction buffer (150 mM Tris-HCl (pH 8), 250 mM NaCl, 25 mM EDTA 0.5 % (w/v) SDS), samples were mixed and centrifuged for 15 min at 13,000 g at room temperature. 150 μL of the supernatant were transferred to a 1.5 mL reaction tube and mixed with 150 μL isopropanol. Precipitated DNA was pelleted by centrifugation at 13,000 g for 15 minutes at room temperature. The DNA pellet was washed with 500 µL of 70 % ethanol and centrifuged for 5 min as above. The supernatant and residual ethanol was removed and the pellet was air-dried for five minutes before being dissolved in 40 µl of TlowE buffer (10 mM Tris-HCl (pH 8), 0.1 mM EDTA).

### Microscopy

For high resolution confocal stacks, six day old seedlings were fixed in a solution containing paraformaldehyde and SCRI Renaissance 2200 (Renaissance Chemicals, North Duffield, UK) as described (Musielak, et al., 2015). After washing in PBS twice, seedlings were transferred to ClearSee (Ursache, et al., 2018; Kurihara, et al., 2015) and additionally stained with 0.2 % basic fuchsin for directly in ClearSee if differentiated xylem was analysed. ClearSee solution was exchanged after a few days. For imaging, samples were in ClearSee in microscopy chambers and stacks of meristems or mature vasculature tissue were acquired with a Leica SP8 with 63x glycerol objective. SCRI Renaissance 2200 fluorescence was excited with the 405 nm laser line, GFP with 488 nm, YFP with 514 nm, emission was collected between 425 and 475 nm (SCRI Renaissance 2200), 520 and 550 nm (GFP) and 530 and 580 nm (YFP). Stacks were processed with the resliced tool in Fiji and segmented at the indicated positions using CellSeT (Pound, et al., 2012).

For pTCSn reporter imaging, longitudinal sections of six day old seedlings root meristems were acquired with a Leica SP5 microscope and a 63x water objective. Venus fluorescence was recorded as described for YFP above, mCherry fluorescence was excited with the 561 nm laser and collected between 575 and 650 nm.

### EdU staining

Plate-grown seedlings were transferred to liquid half-strength MS medium supplemented with 10 µM EdU for the indicated times after which seedlings were either washed in liquid half strength MS twice and incubated in fresh liquid half strength MS without EdU or directly fixed in 4% para-Formaldehyde, 0.1 % Triton X-100 in 1x PBS for 1 hour. After three washes in PBS, the click chemistry reaction was performed with the base-click EdU-Click 488 kit according to the manufacturer’s instructions. The seedlings were washed in PBS three times and transferred to ClearSee solution. Cell walls were stained with Calcofluor white (fluorescent Brightener 29, Sigma-aldrich) in ClearSee according to (Ursache, et al., 2018). Imaging of Alexa488-labelled nuclei was performed as described above for GFP, imaging of Calcofluor white-derived cell wall fluorescence as described above for SCRI Renaissance 2200.

## Supplemental Information titles and legends

**Figure S1. *sgo1* is a recessive mutation**. (A) Schematic representation of the Arabidopsis root meristem (left) and cross sections through the vascular cylinder and the endodermis 15 µm and 150 µm distance from the quiescent centre (QC, right). (B) Protoxylem quantification of Col-0, *sgo1* and segregation of protoxylem phenotype in F2 population derived from a cross between the two genotypes. Graph depicts frequency of roots with the indicated number of protoxylem cells. PT = phenotype.

**Figure S2. Ectopic xylem precursor cells form continuous strands in *sgo1***. pTMO5:GFP-labelled ectopic cell files in a longitudinal cross section (upper panel) and maximum projection of the same confocal stack. Arrowheads indicate two xylem precursor cell files outside the xylem axis.

**Figure S3. *SGO1* encodes AGO10, which is essential for root vascular patterning in Col-0 and L*er***. (A) Basic fuchsin staining of lignified xylem cells in L*er* and *the AGO10* mutant *zll-3*. Upper panels are xy projections of a confocal stack, lower panels optical xy sections through the same stack. Asterisks denote cells with protoxylem differentiation, arrows point to ectopic xylem strands. (B-D) Frequency of roots with the indicated number of protoxylem (B), metaxylem (C) or total xylem (D) cells in L*er* and zll-3. Asterisks indicate statistically significant difference from Col-0 based on Mann-Whitney U test (**P<0.01, *P<0.05). (C) Quantification of vascular cell number in cross section of confocal stacks at 15 µm, 22 µm, and 150 µm distance from the quiescent centre (QC) cells in L*er* and *zll-3* meristems. Graph depicts means ± s.d. and individual data (n = 10-12). Letters in graph indicate statistically significant differences based on Tukey’s post-hoc test after one-way ANOVA. (D) Frequency of roots with the indicated protoxylem cell number in L*er, sgo1* and F1 plants of the indicated crosses, demonstrating that *zll-3* and *sgo1* are allelic to each other. Asterisks indicate statistically significant differences from L*er* based on Dunn’s post hoc test with Benjamini–Hochberg correction after Kruskal–Wallis modified U test (**P <0.01).

**Figure S4. AGO10 is required for HD-ZIP III-mediated vascular patterning**. (A) Exemplary section through a confocal stack of a *pAGO10:GFP* meristem. White bars indicate positions of fluorescence quantification. Magenta trace indicates cell wall fluorescence, green trace fluorescence derived from *pAGO10:GFP*. Note the different y-axes for green and magenta channel. e = endodermis, asterisk = pericycle, m = metaxylem. (B) Quantitative Real Time Reverse Transcribed PCR (pRT-PCR) analysis indicates reduced HD-ZIP III transcript abundance in *sgo1*. Bars indicate average of Mean normalized expression values from three independent biological replicates ± s.d., individual data points are indicated. Asterisks denote statistically significant differences based on a two-tailed student’s ttest of log2-transformed MNE values according to (Rieu and Powers 2009). (C) Root length quantification of L*er, phb-11, phv-13, cna-2*, and *phb phv cna* 7 days after germination. Graph denotes means ±s.d., individual data points are indicated. (D) Quantification of vascular cell number in *35S:STTM165/6* expressing a miR165/6 target mimic, *sgo1*, and the target mimic line in the *sgo1* background. (E) Quantification of mature miR165/6 using a modified stem-loop qPCR (Chen et al., 2005, Dastidar et al., 2019) and pri-MIR166b using qPCR. Asterisks denote statistically significant differences based on a two-tailed student’s ttest of log2-transformed MNE values according to (Rieu and Powers 2009).

**Figure S5. *sgo1* shows enhanced resistance to water-limited conditions**. (A) Meristematic vascular cell number 150 µm from the QC of Col-0 and sgo1 plants seven days after germination on control medium or medium simulating water deficit by addition of 150 g/L PEG8000. Graph indicates mean ± s.d., individual data points are indicated. (B) Root length of plants grown on control medium of medium simulating water deficit by addition of 150 g/L PEG8000 at seven days after germination. Image depicts representative seedlings. Graph indicates mean ± s.d., individual data points are indicated. (C) Root growth 11 days after transfer of five day old seedlings to medium infused with the indicated amount of PEG8000. Graph indicates mean ± s.d., individual data points are indicated.

**Table S1. Oligonucleotides used in this study**

**Table S2. GreenGate constructs**

