## Supplemental Figures for "ARGONAUTE10 is required for cell fate specification and the control of formative cell divisions in the Arabidopsis root meristem"

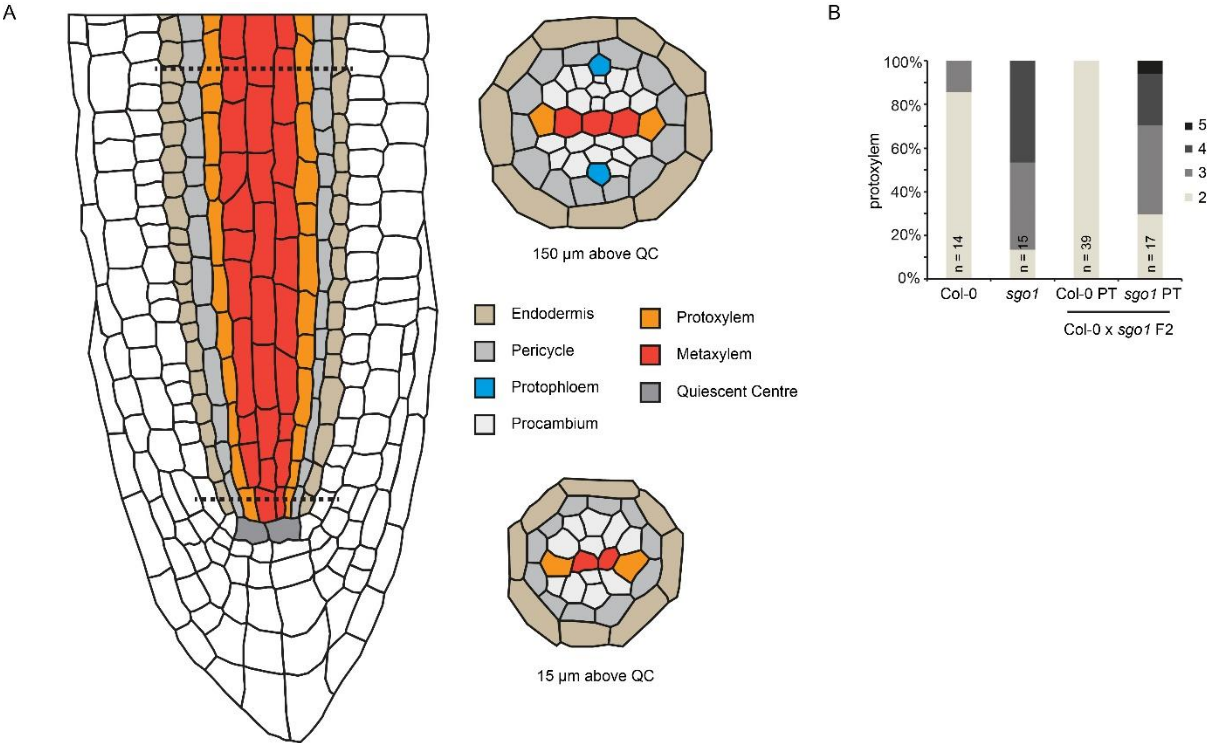


**Figure S1. *sgo1* is a recessive mutation.** (A) Schematic representation of the Arabidopsis root meristem (left) and cross sections through the vascular cylinder and the endodermis 15 µm and 150 µm distance from the quiescent centre (QC, right). (B) Protoxylem quantification of Col-0, *sgo1* and segregation of protoxylem phenotype in F2 population derived from a cross between the two genotypes. Graph depicts frequency of roots with the indicated number of protoxylem cells. PT = phenotype.

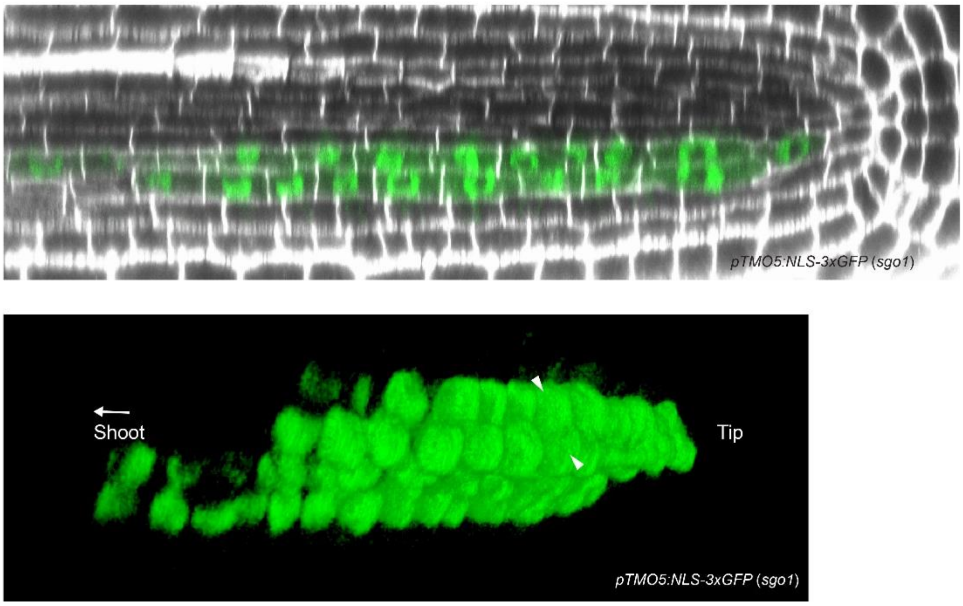


**Figure S2. Ectopic xylem precursor cells form continuous strands in *sgo1*.** pTMO5:GFP-labelled ectopic cell files in a longitudinal cross section (upper panel) and maximum projection of the same confocal stack. Arrowheads indicate two xylem precursor cell files outside the xylem axis.

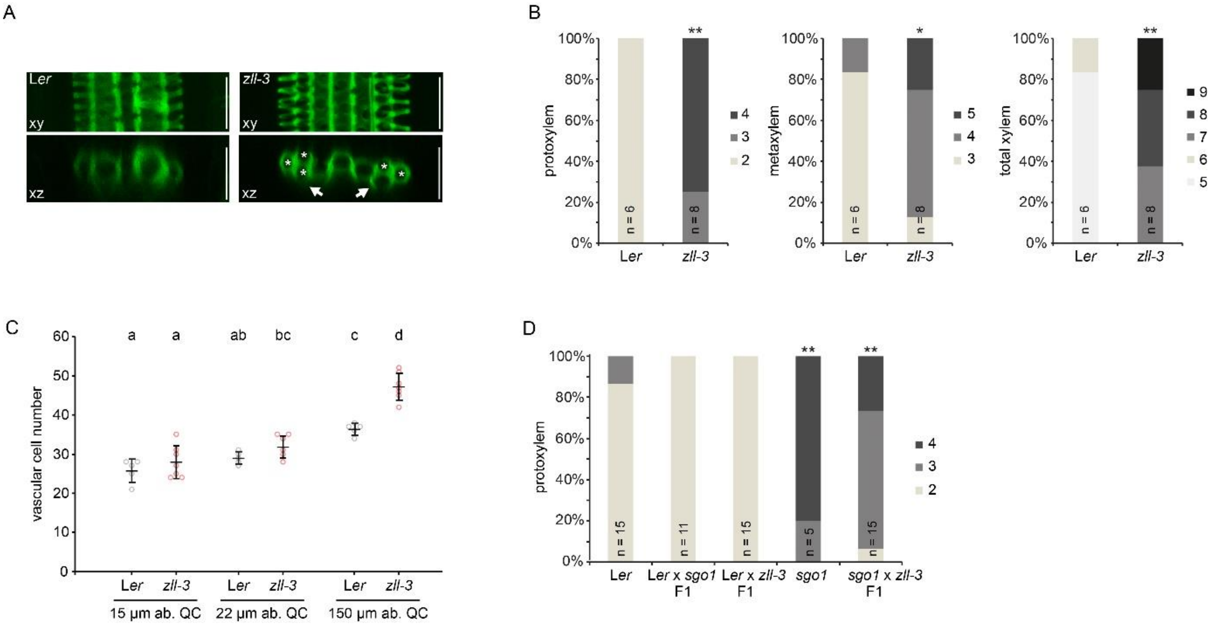


**Figure S3. *SGO1* encodes AGO10, which is essential for root vascular patterning in Col-0 and L*er*.** (A) Basic fuchsin staining of lignified xylem cells in L*er* and *the AGO10* mutant *zll-3*. Upper panels are xy projections of a confocal stack, lower panels optical xy sections through the same stack. Asterisks denote cells with protoxylem differentiation, arrows point to ectopic xylem strands. (B-D) Frequency of roots with the indicated number of protoxylem (B), metaxylem (C) or total xylem (D) cells in L*er* and zll-3. Asterisks indicate statistically significant difference from Col-0 based on Mann-Whitney U test (**P<0.01, *P<0.05). (C) Quantification of vascular cell number in cross section of confocal stacks at 15 µm, 22 µm, and 150 µm distance from the quiescent centre (QC) cells in L*er* and *zll-3* meristems. Graph depicts means ± s.d. and individual data (n = 10-12). Letters in graph indicate statistically significant differences based on Tukey's post-hoc test after one-way ANOVA. (D) Frequency of roots with the indicated protoxylem cell number in L*er*, *sgo1* and F1 plants of the indicated crosses, demonstrating that *zll-3* and *sgo1* are allelic to each other. Asterisks indicate statistically significant differences from L*er* based on Dunn's post hoc test with Benjamini–Hochberg correction after Kruskal–Wallis modified U test (**P <0.01).

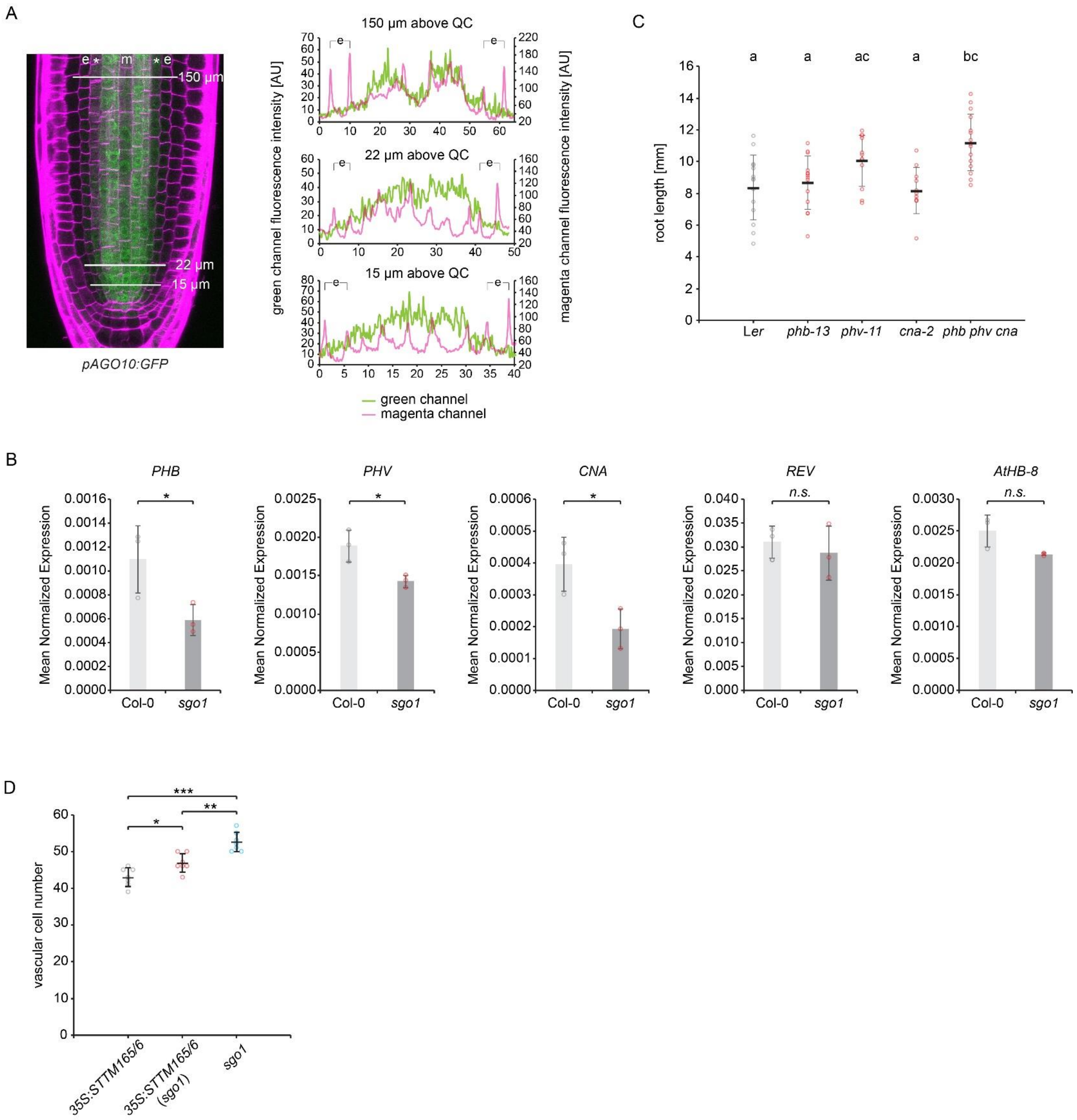


**Figure S4. AGO10 is required for HD-ZIP III-mediated vascular patterning.** (A) Exemplary section through a confocal stack of a *pAGO10:GFP* meristem. White bars indicate positions of fluorescence quantification. Magenta trace indicates cell wall fluorescence, green trace fluorescence derived from *pAGO10:GFP*. Note the different y-axes for green and magenta channel. e = endodermis, asterisk = pericycle, m = metaxylem. (B) Quantitative Real Time Reverse Transcribed PCR (pRT-PCR) analysis indicates reduced HD-ZIP III transcript abundance in *sgo1*. Bars indicate average of Mean normalized expression values from three independent biological replicates ± s.d., individual data points are indicated. Asterisks denote statistically significant differences based on a two-tailed student's ttest of log2-transformed MNE values according to (Rieu and Powers 2009). (C) Root length quantification of L*er*, *phb-11*, *phv-13*, *cna-2*, and *phb phv cna* 7 days after germination. Graph denotes means ±s.d., individual data points are indicated. (D) Quantification of vascular cell number in *35S:STTM165/6* expressing a miR165/6 target mimic, *sgo1*, and the target mimic line in the *sgo1* background.

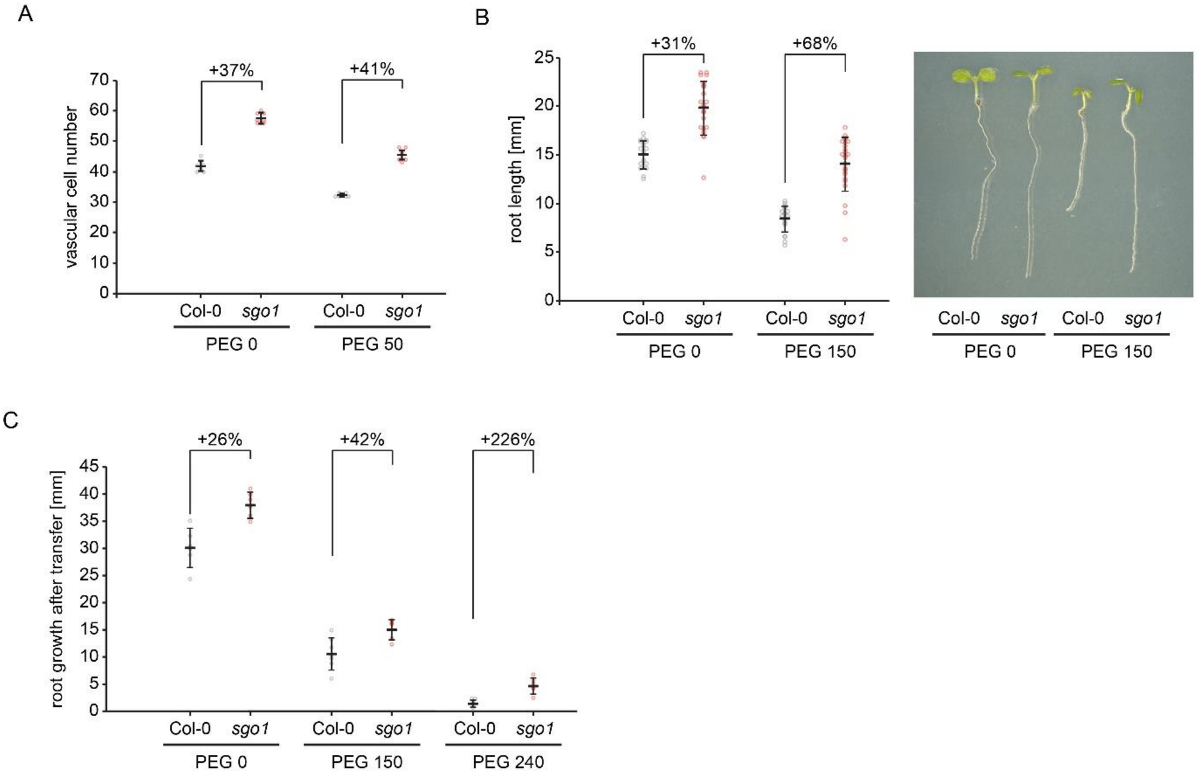


**Figure S5. *sgo1* shows enhanced resistance to water-limited conditions.** (A) Meristematic vascular cell number 150 µm from the QC of Col-0 and sgo1 plants seven days after germination on control medium or medium simulating water deficit by addition of 150 g/L PEG8000. Graph indicates mean ± s.d., individual data points are indicated. (B) Root length of plants grown on control medium of medium simulating water deficit by addition of 150 g/L PEG8000 at seven days after germination. Image depicts representative seedlings. Graph indicates mean ± s.d., individual data points are indicated. (C) Root growth 11 days after transfer of five day old seedlings to medium infused with the indicated amount of PEG8000. Graph indicates mean ± s.d., individual data points are indicated.
