## Supplementary material for "ARGONAUTE10 is required for cell fate specification and the control of formative cell divisions in the Arabidopsis root meristem": Table S1

*Table S1. Oligonucleotides used in this study.*

| Oligo name | Sequence | Purpose |
|---|---|---|
| CER448567_F | ATA GAA AGG TTT GAG GGG GC | Bulked segregant analysis |
| CER448567_R | TGC GAA GAA CCA CTA AAC CC | Bulked segregant analysis |
| F9L1_F | CTC GGA AAT TCT TAG CTT TC | Bulked segregant analysis |
| F9L1_R | TTA TAA CTT GCC CAA AGC GAA | Bulked segregant analysis |
| F1K23ind38_F | GGA TTG AAC ATA GGG AAG GGG | Bulked segregant analysis |
| F1K23ind38_R | GAT CTG TAT CTG AAA CCT GGG | Bulked segregant analysis |
| CER464787-Indel-44_F | TTT GAA CTA ACC TTC TGA GG | Bulked segregant analysis |
| CER464787-Indel-44_R | CAT GTT GAT GAT TCA ATT GC | Bulked segregant analysis |
| F6D8ind94_F | CCG TTA CCC CCA TAC GAA CG | Bulked segregant analysis |
| F6D8ind94_R | TCG TGA GGT TAT GCC GAT CC | Bulked segregant analysis |
| F5I14_F | CTG CCT GAA ATT GTC GAA AC | Bulked segregant analysis |
| F5I14_R | GGC ATC ACA GTT CTG ATT CC | Bulked segregant analysis |
| CER459153_F | TCG TGA CCA AAT CCT GAA CA | Bulked segregant analysis |
| CER459153_R | TGT CCA AGT AAT GCC GTG AG | Bulked segregant analysis |
| CER466780_F | GAA CCC TTA TAA TAT GGC TGG C | Bulked segregant analysis |
| CER466780_R | GGA AGT ATT CCC AAG ACA AGG | Bulked segregant analysis |
| MSAT2-36_F | GAT CTG CCT CTT GAT CAG C | Bulked segregant analysis |
| MSAT2-36_R | CCA AGA ACT CAA AAC CGT T | Bulked segregant analysis |
| F3N11_F | GTT AAA GCG AGG ACG ATT GG | Bulked segregant analysis |
| F3N11_R | AGA TAC TGT CGC CAT CAA GG | Bulked segregant analysis |
| T2P4_F | ACT AGT CCC ACT GTC GAT C | Bulked segregant analysis |
| T2P4_R | GTT ACT TCG TAA GTC CCT AC | Bulked segregant analysis |
| MSAT2-9_F | TAA AAG AGT CCC TCG TAA AG | Bulked segregant analysis |
| MSAT2-9_R | GTT GTT GTT GTG GCA TT | Bulked segregant analysis |
| nga172_F | AGC TGC TTC CTT ATA GCG TCC | Bulked segregant analysis |
| nga172_R | CAT CCG AAT GCC ATT GTT C | Bulked segregant analysis |
| CER455386_F | CTC TTT TGG CTC GGA CAA G | Bulked segregant analysis |
| CER455386_R | GTT GTA ATC GGG AAA ATG C | Bulked segregant analysis |
| CER455914_F | GGA GCA GAG AAA GAG AC | Bulked segregant analysis |
| CER455914_R | GAG GAA GGA CAA CAT GGC | Bulked segregant analysis |
| CER456071-Indel-35_F | AGC CAT AGG TAA TGT CCA CG | Bulked segregant analysis |

| | | |
|---|---|---|
| CER456071-Indel-35_R | CTC GCG GAT GAG TAT CAT CC | Bulked segregant analysis |
| CER470441_F | GCT AAC AGG GAT ATC AAA TGT GC | Bulked segregant analysis |
| CER470441_R | CGG ACG AGC TGA CAC TTG TA | Bulked segregant analysis |
| CER470172_F | GTA AAA CTC CTC CTC TGG GG | Bulked segregant analysis |
| CER470172_R | TGT AAT CGT GGC GGA ACG GG | Bulked segregant analysis |
| CER459609_F | TCG CTT TTG AAG ATT TGT GC | Bulked segregant analysis |
| CER459609_R | GGG AGC TTC TCA GTG GTC TG | Bulked segregant analysis |
| nga8_F | GAG GGC AAA TCT TTA TTT CGG | Bulked segregant analysis |
| nga8_R | TGG CTT TCG TTT ATA AAC ATC C | Bulked segregant analysis |
| FCA0ind25_F | AAG CCA ACT ATT GCC AAG GG | Bulked segregant analysis |
| FCA0ind25_R | TCA CTG CCC TTT ACT CCG GT | Bulked segregant analysis |
| F7J7-47_F | TGG TGA AGA GCT TAG TTG ATG A | Bulked segregant analysis |
| F7J7-47_R | TCA CTA GAT ATC TCT AGT GGC T | Bulked segregant analysis |
| CER451534_F | AGC TAC GGT GGA GTG TAA TTT CGT | Bulked segregant analysis |
| CER451534_R | GCT GAT ACT TGC TTT CGC TTT GCA G | Bulked segregant analysis |
| CER459444_F | AGT AGC ATC GTA GCT CCT AGG | Bulked segregant analysis |
| CER459444_R | GTT GTA TAC GTG CAC GTT CCC | Bulked segregant analysis |
| CER456519_F | TGC TAA AAT ATA AAA CTT CC | Bulked segregant analysis |
| CER456519_R | TTA TGC AGA TGT ATG AGG CC | Bulked segregant analysis |
| nga151_F | GTT TTG GGA AGT TTT GCT GG | Bulked segregant analysis |
| nga151_R | CAG TCT AAA AGC GAG AGT ATG ATG | Bulked segregant analysis |
| nga139_F | GGT TTC GTT TCA CTA TCC AGG | Bulked segregant analysis |
| nga139_R | AGA GCT ACC AGA TCC GAT GG | Bulked segregant analysis |
| T26D22-IND52/CER459812_F | TCC CAC GAA GAG AGA AGT GC | Bulked segregant analysis |
| T26D22-IND52/CER459812_R | CTA TTT GCT TAT GAA GGT GTC C | Bulked segregant analysis |
| CER456772_F | CCA TGT GAC ATG CAC TTA CAC | Bulked segregant analysis |
| CER456772_R | ACC ATT CTC TAC CAC TCC AC | Bulked segregant analysis |
| K6M13ind33/CER454758_F | ATA GAT GAG ATC CAC TTG CC | Bulked segregant analysis |
| K6M13ind33/CER454758_R | ACA AAC TGT TGC TGT GGG AG | Bulked segregant analysis |
| MBK5ind35/CER455203_F | ATT CTC GGA CCA GGC TTC AT | Bulked segregant analysis |
| MBK5ind35/CER455203_R | AAA GAA CAG CTA CTG CGT GC | Bulked segregant analysis |
| nga151_R | CAG TCT AAA AGC GAG AGT ATG ATG | Bulked segregant analysis |

| | | |
|---|---|---|
| nga139_F | GGT TTC GTT TCA CTA TCC AGG | Bulked segregant analysis |
| nga139_R | AGA GCT ACC AGA TCC GAT GG | Bulked segregant analysis |
| T26D22-IND52/CER459812_F | TCC CAC GAA GAG AGA AGT GC | Bulked segregant analysis |
| T26D22-IND52/CER459812_R | CTA TTT GCT TAT GAA GGT GTC C | Bulked segregant analysis |
| At5g43810_sgo1_dCAPS_F | GTTTCCAGTAGATGAGAACTG | dCAPS marker (HindIII) |
| At5g43810_sgo1_dCAPS_R | AACAAACAAACCAGCAGTTTACGGTTAAGCT | dCAPS marker (HindIII) |
| At5g47720_Col_dCAPS_F | CTGGGATTAGATCCTgtaagtc | dCAPS marker (HinfI) |
| At5g47720_Col_dCAPS_R | CCCAATGGATGTCCCAGTGAAACAGCGACT | dCAPS marker (HinfI) |
| At5g51560_CAPS_F | ctctcgaggcaaatgtgtaat | CAPS marker (XmnIII) |
| At5g51560_CAPS_R | AGGGTTGAAGAAATGACATCC | CAPS marker (XmnI) |
| qPHB-F | CTTTGGTAGTGGCGTGCTTT | qRT-PCR |
| qPHB-R | GCCCATTCAGATCGGTGTTC | qRT-PCR |
| qPHV-F | CCAAGATCATGCAGCAGGGA | qRT-PCR |
| qPHV-R | CGCTTGCTCATACGAAACCG | qRT-PCR |
| qCNA-F | AGAATGTTCCTCCGGCGATC | qRT-PCR |
| qCNA-R | TGCCCTCCAAATCCTCCAAC | qRT-PCR |
| qPri-MIR166b-F | TGTCTGGCTCGAGGACTCTT | qRT-PCR |
| qPri-MIR166b-R | TCCGACGACACTAAAACCCT | qRT-PCR |
| qmiR166-F | TCGCTTCGGACCAGGCTTCA | qRT-PCR |
| qmiR166-R | GTGCAGGGTCCGAGGT | qRT-PCR |
| miR166stem-loop | GTCGTATCCAGTGCAGGGTCCGAGG TATTCGCACTGGATACGACGGGGAA | Stem-loop RT primer |
| ACT2-F | CTAAGCTCTCAAGATCAAAGGCTTA | qRT-PCR |
| ACT2-R | ACTAAAACGCAAAACGAAAGCGGTT | qRT-PCR |
| A04555 | AACAGGTCTCCGGCTAAAAGCTTGACTAGT CAAAGATC | Cloning of pGGA043 (pTCSn) |
| A04630 | AAC AGG TCT CaT GTT TCT CCA AAT GAA ATG AAC TTC | Cloning of pGGA043 (pTCSn) |
| A03821 | AAC AGG TCT CAA ACA ATG GTG AGC AAG GGC GAG | Cloning of pGGB007 |
| A02228 | AAT TCG GTC TCA AGC CAG CAA TTG CTG CGG CAG C | Cloning of pGGB007 |
| A02530 | AAC AGG TCT CAG GCT CCA TGG TGA GCA AGG GCG AGG A | Cloning of pGGC023 |
| A02531 | AAC AGG TCT CAC TGA CTT GTA CAG CTC GTC CAT G | Cloning of pGGC023 |
