## Supplementary material for "ARGONAUTE10 is required for cell fate specification and the control of formative cell divisions in the Arabidopsis root meristem": Table S2

**Table S3**. Overview of constructs generated with GreenGate cloning (Lampropoulos et al., 2013) and the primers used (see oligo table for sequences) to generate modules, where appropriate.

| | | | |
|---|---|---|---|
| **pCW178** | **pTCSn:2xVenus-NLS** | | |
| pGGA043 | pTCSn | A04555 | A04630 |
| pGGB007 | Venus-linker | A03821 | A02228 |
| pGGC023 | Venus | A02530 | A02531 |
| pGGD007 | Linker-NLS | Lampropoulos et al., 2013 | |
| pGGE001 | *RBCS* terminator | Lampropoulos et al., 2013 | |
| pGGF001 | pMAS:BastaR:tMAS | Lampropoulos et al., 2013 | |
| pGGZ0003 | destination vector | Lampropoulos et al., 2013 | |
| **pCW066** | **pUBQ10:3xmCherry-NLS** | | |
| pGGA006 | UBQ10 (At4g05320)promoter | Lampropoulos et al., 2013 | |
| pGGB003 | B-Dummy | Lampropoulos et al., 2013 | |
| pGGC026 | 3xmCherry | Lampropoulos et al., 2013 | |
| pGGD007 | Linker-NLS | Lampropoulos et al., 2013 | |
| pGGE009 | UBQ10 terminator | Lampropoulos et al., 2013 | |
| pGGF007 | pNOS:KanR:tNOS | Lampropoulos et al., 2013 | |
| pGGZ003 | destination vector | Lampropoulos et al., 2013 | |
